# Rcirc: an R package for circRNA analyses and visualization

**DOI:** 10.1101/2020.03.06.980151

**Authors:** Peisen Sun, Haoming Wang, Guanglin Li

**Affiliations:** Key Laboratory of Ministry of Education for Medicinal Plant Resource and Natural Pharmaceutical Chemistry, Shaanxi Normal University, Xi’an, China; College of Life Sciences, Shaanxi Normal University, Xi’an, China; College of Plant Protection, Northwest A&F University, Yangling, China

**Keywords:** Bioinformatics, circular RNAs, visualization, R package, coding potential, Ribosome profiling

## Abstract

Circular RNA (circRNA), which has a closed-loop structure, is a kind of special endogenous RNA and plays important roles in many biological processes. With the improvement of next-generation sequencing technology and bioinformatics methods, some tools have been published for circRNA detection based on RNA-seq. However, only a few tools focus on downstream analyses, and they have poor visualization ability. Here, we developed the R package ‘Rcirc’ for further analyses of circRNA after its detection. Rcirc identifies the coding ability of circRNA and visualize various aspects of this feature. It also provides general visualization for both single circRNAs and meta-features of thousands of circRNAs. Rcirc was designed as a user-friendly tool that covers many highly automatic functions without running many complicated processes by users. It is available on GitHub (https://github.com/PSSUN/Rcirc) under the license GPL 3.0.

## Introduction

Circular RNA (circRNA) is an abundant functional RNA molecule with a highly conserved closed-loop structure that is generated by back-splicing without 5’ cap and 3’ poly (A) tails(Vicens and Westhof, 2014). Single-stranded circRNAs appear to play a role in endogenous cells, such as immune regulation and cancer and RNA-binding protein regulation(Li et al., 2018). In the medical field, circRNAs are often used as biomarkers to identify the occurrence of cancer because of their ring structure, which is difficult for RNase R to degrade in the liquid phase(Bonizzato et al., 2016)(Kulcheski et al., 2016).

In addition, circRNA has translation capabilities and important biological significance(Pamudurti et al., 2017). Organisms contain a large number of short peptides (<100 aa) that play an important role in many regulatory pathways(Delcourt et al., 2018)(Slavoff et al., 2013). Recently, many studies have shown multiple pieces of evidence that strongly support circRNA translation, such as the specific association of circRNAs with translating ribosomes(Pamudurti et al., 2017). This means that a large number of circRNAs also have a small open reading frame (ORF) that can be translated and produce functional peptides. However, it is difficult for the traditional RNA-seq technique to accurately identify these short peptides from transcripts. In recent years, the increasing maturity of ribosome profiling technology, which provides strong evidence for translation events, has enabled accurate identification of short peptides. In contrast to the traditional RNA-seq technique, ribosome profiling can confirm the specific translation region of mRNAs(Brar and Weissman, 2015).

Feature analysis is also a very important area in the field of circRNA, and most published circRNAs have been analyzed. Especially in the field of machine learning, the existence of numerous machine learning studies for identifying new circRNAs has left no doubt that a large number of possible features are needed, including the types of shear signal, the length of circRNAs, the frequency of triplet codons, and the location of different back-splice junctions on the genome, but there is currently no tool for characterizing circRNAs and subsequent visualization.

Based on high-throughput sequencing data, various tools have been developed for circRNA detection, including circRNA finder(Westholm et al., 2015), find_circ(Memczak et al., 2013), CIRCexplorer(Zhang et al., 2014), and CIRI(Gao et al., 2015). Additionally, numerous databases for circRNA that are based on those detection tools have also been published(Hansen et al., 2016).

However, most published software programs are designed to predict circRNAs, and only two software programs can identify their coding capability(Meng et al., 2017)(Sun and Li, 2019). There is no tool that focuses on downstream analyses and visualization of features or mapping for circRNAs after the prediction process. Circtools can help design primers but cannot perform large-scale analysis and visualize the read mapping(Jakobi et al., 2018). Here, we developed Rcirc, an R package. It can not only identify candidate circRNAs but also recognize the translation ability of circRNAs. Rcirc also performs many feature analyses and data visualization. Through a display of diversity, users can easily see the various sequence features of a certain data set and determine whether a feature is representative.

## Implementation

Currently, Rcirc contains 10 functions for the analysis of circRNAs. Rcirc covers three main parts for circRNA research (Figure 1): circRNA detection, coding ability identification and feature visualization. The users can run any functions from those parts individually or run all functions of the whole pipeline.

**Figure 1.**
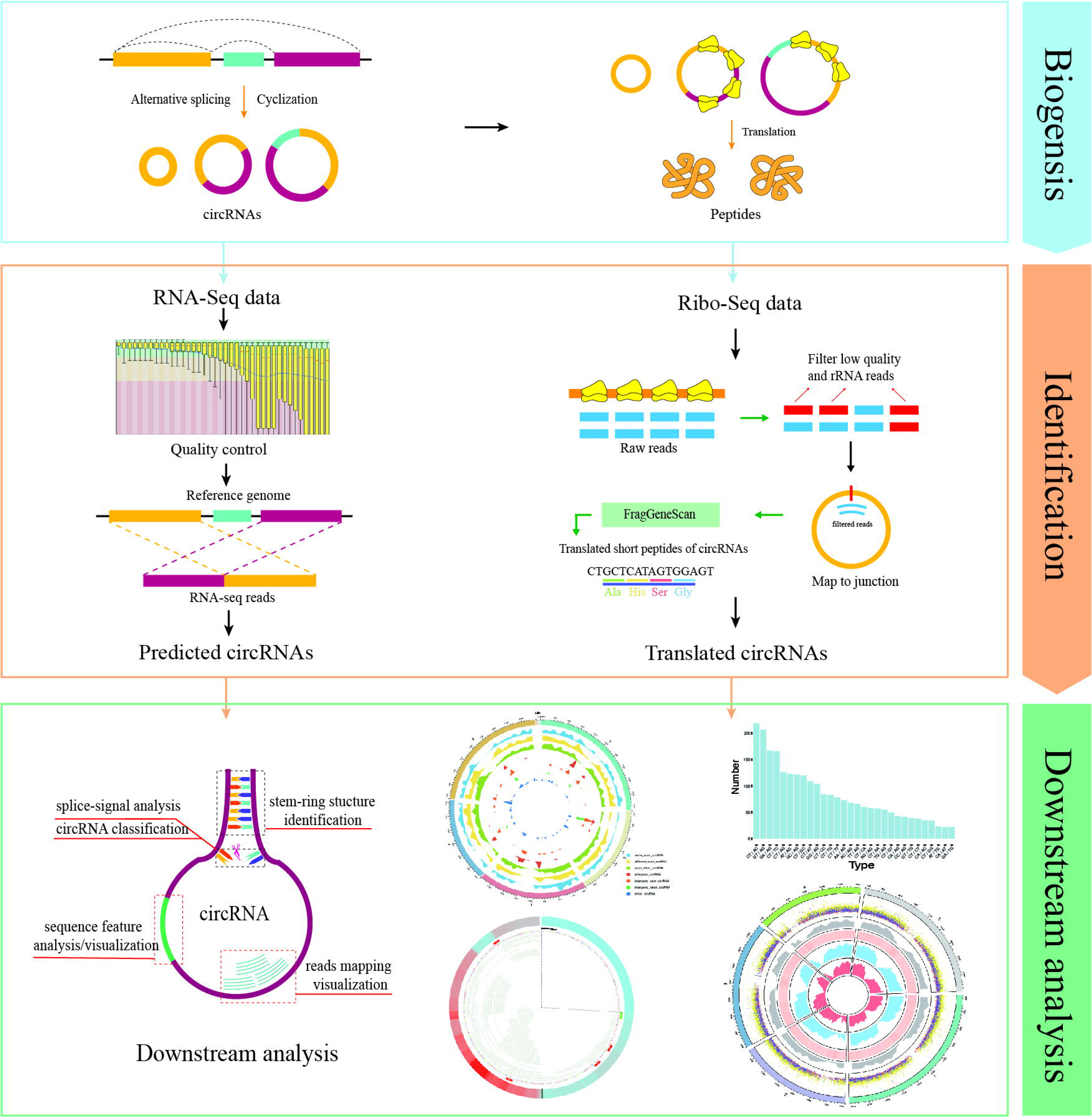
Rcirc workflow. From top to bottom are shown the identification of circRNAs, identification of circRNA translation ability, and downstream analysis/visualization of circRNA. The right column represents the analyses that Rcirc can perform at the corresponding stage.

### CircRNA detection and coding ability identification

Users can make a de novo prediction for circRNAs based on RNA-seq data by the function *predictCirc*, which aids in fundamental quality control for RNA-seq data and circRNA prediction by calling CIRI2. Finally, a predicted fasta file and prediction report in csv format are outputted to an appointed file.

The function *translateCirc* can help to identify the coding capability from the given circRNAs based on Ribo-seq data. Because the ribosome needs to span the back-splice junction composed of the 3’ end and the 5’ end during translation, if a circRNA has translational behavior, the Ribo-seq reads can be mapped to the back-splice junction, which provides the criterion for the translation of circRNA(Sun and Li, 2019).

Since the sequence spanning the back-splice junction in circRNA is spliced from the 3’ and 5’ ends, it cannot be directly obtained. Therefore, in Rcirc, we completely copy each circRNA sequence and concatenate it in the original sequence. Later, the linear sequence was used to simulate the real situation of the circRNA at the back-splice junction. After that, the reads of the ribosomal maps are aligned to this linear sequence.

Compared to traditional RNA-seq data, ribosome profiling data require further processing to remove the rRNA sequence in addition for the routine quality control process (removing the linker, filtering the low-quality fragments) because the fragments of rRNA during sequencing may also be mixed in the final data, causing interference with the results. Since the length of the reads obtained by the ribosome data is relatively short (generally less than 50 bp), even if there is a successful comparison of the reads to the back-splice junction, it is not enough to indicate that the reads are from this position because they may also originate from any similar area on the transcriptome.

To avoid this issue, Rcirc first aligns all processed Ribo-seq reads back to the rRNA sequence and reference genome, removing all reads that can be aligned to the rRNA and linear transcripts and leaving only reads that were not successfully aligned. This step ensures that all remaining reads have no similar regions on the rRNA and the linear transcript. After this, Rcirc aligns these reads to the previously simulated back-splice junction to see if it matches and then counts the number of reads that can align to the back-splice junction. The circRNA is finally identified as a translated circRNA if there are no less than 3 Ribo-seq reads on its back-splice junction.

### Downstream analysis and visualization

In this section, we introduce 4 commonly used functions.

The *mappingPlot* function is one of the important functions in Rcirc. Most next-generation sequencing (NGS) data visualization browser tools, such as IGV, help to view the mapping between reads and sequences. This method of expression is more vivid and intuitive than files in text formats such as the SAM/BAM format and thus helps researchers better study the problem in the interval of interest. However, it is impossible for these tools to view mapping results on circRNA because of its end-to-end ring structure, which is different from linear structures. With the *makeGenome* function, this issue has been solved. It automatically connects the 5’ end and 3’ end of circRNA as a ring and produces a data frame in R that contains all the mapping information of each circRNA. In this function, the mapping of Ribo-seq data on each junction can be revealed clearly by a ring diagram, which simulates the real circular form of circRNA in cells. This visualization includes reads covered region, reads covered density, highlighted bases, start codons and stop codons. Moreover, users can be free to enlarge the mapping region by an optional parameter. The detailed usage of *mappingPlot* can be found in Rcirc user documents.

circRNAs are generally classified by the location of the back-splice junction. Here, we use *classByType* to classify a given circRNA. According to the position of the back-splice junction, we divide the circRNA into 7 categories: *same_exon*, *different_exon*, *intron_exon*, *intron*, *intron_intergenic*, *exon_intergenic* and *intergenic*. After completing the classification, *classByType* can give a classification table and a ring density map of the distribution of different types of circRNAs on different chromosomes. A specific demonstration can be seen in Figure 2 (B). Users need to enter only an annotation file of the genome and a BED format file of circRNAs, which can be easily analyzed.

**Figure 2.**
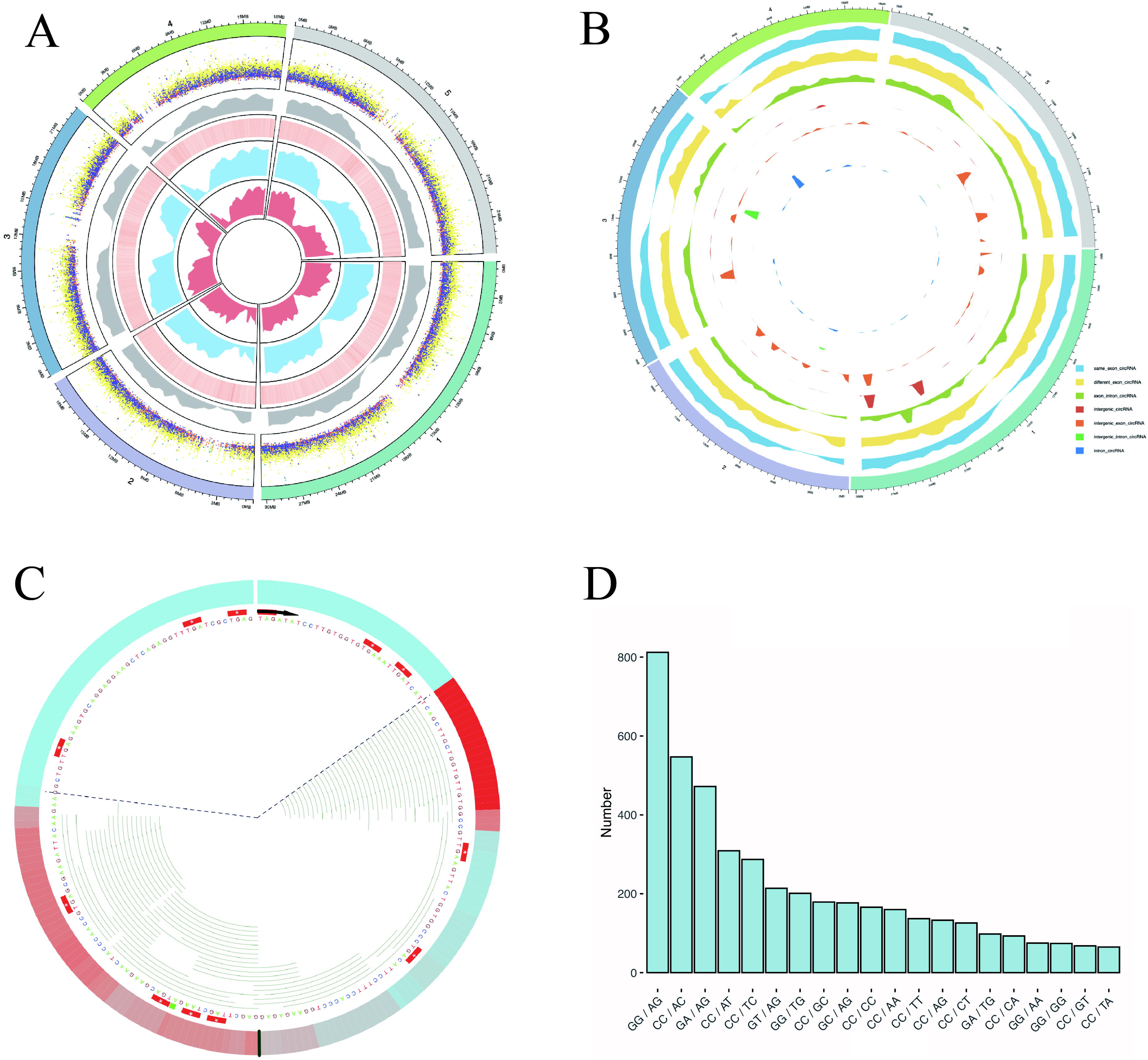
Some characterizations of *Arabidopsis thaliana* were performed using Rcirc. **(A)** An overview of the characteristics of all candidate circRNAs in *A. thaliana*. From the outside to the inside is shown a distribution of different types of circRNAs of different lengths in dot plot (the height of the dots represents the length of the circRNA, different colors represent different types of circRNA, orange, green, light blue, cyan, dark blue, purple, light yellow represent: same_exon_circ, diff_exon_circ, exon_intron_circ, intron_circ, interg_circ, interg_exon_circ, interg_intron_circ, respectively); a density distribution of all the circRNAs; a heatmap of GC content for the genome, wherein red reveals the high GC content regions and white reveals the low GC content regions; the RNA-seq read coverage density (blue); and the Ribo-seq read coverage density (red). **(B)** Classification of circRNAs and plot density profiles. Different colors represent different types of circRNAs. **(C)** A visual analysis of the Ribo-seq reads mapping of one of the circRNAs. The black line represents the back-splice junction. The color change in the outermost circle represents the coverage density of the reads at the site. The redder the color is, the greater the coverage density, and the bluer the color is, the smaller the coverage density. Each green line inside represents a Ribo-seq read. The area within the dashed line represents the full length covered by the reads, and the area outside of the dashed line represents the area without read coverage. **(D)** Splice signals of all circRNAs were analyzed by Rcirc, and the resulting histograms are shown.

The *stemRing* function can help to determine the possible stem-ring structure for the given circRNAs in BED format. It extracts the sequence upstream and downstream of each circRNA and makes a local alignment between the downstream sequence and the reverse complement of the sequence upstream. Finally, stemRing outputs the result in a csv format file that contains the position information of each circRNA and the local upstream and downstream alignment information. Using *stemRing* and looking at the results, it is possible to conduct a large-scale investigation of circRNA stem-ring structures and construct a corresponding expression vector containing reverse complementary paired sequences to help circRNA circularize in cells.

The *showOverview* function provides an overview of all circRNAs, including a large amount of information, in one circle. It includes the distribution of circRNAs of different lengths on different chromosomes and the density distribution of circRNAs on different chromosomes. The high-GC and low-GC regions of the genome are labeled with different colors, allowing users to find connections between different features.

In addition, Rcirc includes many other functions. For example, it can also perform joint analysis on thousands of circRNAs to analyze and visualize their distribution as a function of lengths, type, and splice signal.

### Software construction

Rcirc is an R-based toolkit, and all code is written in R language. In the prediction phase, we predict the circRNA by calling the external program CIRI2. The main program of CIRI2 is already included in Rcirc without the user having to download and install it. In the identification phase of translation capabilities, we call STAR, bowtie and trimmomatic to filter and align Ribo-seq data. In the analysis and visualization section, the R packages we use are circlize(Gu et al., 2014), ggplot2(Hadley, 2016), Biostrings(H et al., 2019), IRanges(Lawrence et al., 2013), and others.

Most of the features and their introductions included in Rcirc are shown in Table 1. More details are provided in the Rcirc user manual.

**Table 1.** This table shows all the features of Rcirc. The left column is the name of the function, and the right column is a brief introduction to the function.

| Function | Description |
| --- | --- |
| *makeGenome* | Stitching candidate circRNAs into virtual genomes |
| *PredictCirc* | Identify new circRNA |
| *TranslateCirc* | Identify the translation capabilities of circRNA |
| *showOverview* | Analyze the characteristics of all candidate circRNAs and then render the image |
| *mappingPlot* | Draw an image of the reading mapping near the circRNA back-splice junction |
| *mappingTable* | Combine the Ribo-seq mapping of all candidate circRNAs into a table |
| *stemRing* | circRNA stem-ring structure recognition |
| *classByType* | Classify all circRNAs according to the position of the back-splice junction and draw an image |
| *showJunction* | Analyze the splice signals of all candidate circRNAs and plot charts |
| *showLength* | Show the distribution of the length of circRNAs |
| *showDistribution* | Show the distribution of circRNA on different chromosomes |
| *downloadCircRNA* | Download circRNA from known databases |

## Result and discussion

To demonstrate the Rcirc analysis process, we predicted circRNAs in *Arabidopsis thaliana* using Rcirc, analyzed these circRNAs and visualized the results. The results of the partial analysis are shown in Figure 2.

Most of the currently published software is used to predict circRNA(Hansen et al., 2016), and no software for characterization of circRNA is available. Similar software such as circtools provides partial analysis and visualization functions such as primer design, but its visualization of the back-splice junction remains only for the linear structure, which is less intuitive than the circular structure (Table 2). Another important feature of Rcirc is to fill the gap in this field. In the circRNA literature, many articles have analyzed the characteristics of circRNA from multiple angles, generally including its length distribution, chromosome distribution, classification and shearing according to the location of its back-splice junction in linear transcripts, the shear signal distribution of the back-splice junction, etc. We summarize these analyses and add as much of the downstream analysis as possible in Rcirc. Software with complicated use requires that the user spend much time on learning costs. In Rcirc, we simplified all the analysis processes as much as possible. To perform the above analysis, we do not need to write complex code. The design principle of Rcirc is to complete one analysis using only one line of code. Therefore, Rcirc is an easy-to-use R package for circRNA investigation.

**Table 2.** Comparison of Rcirc and circtools

| Content | Rcirc | circtools |
| --- | --- | --- |
| Frame | R | Python 3 and R |
| Graphical user interface | No | No |
| circRNA detection | CIRI2 | DCC |
| Show the mapping quality | No | Yes |
| Coding ability identify | Yes | No |
| Visualization of mapping region | Yes | No |
| Variation in circRNAs | No | Yes |
| Stem-ring structure recognition | Yes | No |
| Meta-feature analysis | Yes | No |
| Identity individual exons | No | Yes |
| Primer design | No | Yes |

To simplify Rcirc use for feature investigation and visualization, we eliminated a large number of possible input parameters and made the functions highly modular. The user only needs to enter the necessary file path to obtain the final analysis results. This design reduces the use threshold of Rcirc, allowing the user to execute the desired analysis without requiring much cost for learning. However, due to the lack of customizability of the analysis results, the analysis process cannot be directly defined by modifying the parameters. To improve the customizability while simplifying the use as much as possible, we have added some parameters for modifying the way in which the results are displayed, and they are all set to default values for the user to call when needed.

## Conclusion

With the deepening of research on circRNA, an increasing number of studies have proven that it plays an important role in the body. However, the corresponding analysis tools for circRNA have not been developed. The main goal of our study was to develop a user-friendly tool that covers the main demands for circRNA research. Rcirc is a capable and user-friendly package based on the R language. The package provides numerous analyses for both upstream and downstream research, including circRNA detection, coding ability identification, single feature analyses and visualization of meta-features. Furthermore, the users can visualize the read mapping for each back-splice junction of circRNA by using Rcirc with sequencing data. With growing attention on circRNA, Rcirc will become an auxiliary tool to encourage researchers to proceed with further analyses on circRNA, and we will add the most common features into Rcirc in future releases. All the details of usage are included in the Rcirc documents in the GitHub online pages.

## Availability and requirements

Rcirc is available at https://github.com/PSSUN/Rcirc; operating system(s): Linux; programming language: R; other requirements: bowtie, STAR, R packages (circlize, ggplot2, Biostrings, GenomicAlignments, GenomicFeatures, GenomicRanges, IRanges). The installation packages for all of the required software are available on the Rcirc homepage. Users do not need to download the required software individually. The Rcirc home page also provides detailed user manuals for reference. The tool is freely available. There are no restrictions to use by nonacademics.

## Authors’ contributions

SP and WH developed the software package under the guidance of LG, and SP performed all analyses in the manuscript. SP and LG drafted and revised the manuscript. All the authors read and approved the final manuscript.

